# Selective consistency of recurrent neural networks induced by plasticity as a mechanism of unsupervised perceptual learning

**DOI:** 10.1101/2023.06.30.547192

**Authors:** Yujin Goto, Keiichi Kitajo

## Abstract

Understanding the mechanism by which the brain overcomes its inherent inconsistency in activity to achieve consistent information processing is one of the major challenges in neuroscience. Recently, it has been reported that the consistency of neural responses to stimuli that are presented repeatedly is enhanced implicitly in an unsupervised way, and results in improved perceptual consistency. Here, we propose the term "selective consistency" to describe this input-dependent consistency and hypothesize that it will be acquired in a self-organizing manner by plasticity within the neural system. To test this, we investigated whether a reservoir-based plastic model could acquire selective consistency to repeated stimuli. We used white noise sequences randomly generated in each trial and referenced white noise sequences presented multiple times. The results showed that the plastic network was capable of acquiring selective consistency rapidly, with as little as five exposures to stimuli, even for white noise. The acquisition of selective consistency could occur independently of performance optimization, as the network’s time-series prediction accuracy for referenced stimuli did not improve with repeated exposure and optimization. Furthermore, the network could only achieve selective consistency when in the region between order and chaos. These findings suggest that the neural system can acquire selective consistency in a self-organizing manner and that this may serve as a mechanism for certain types of learning.

## 1. Introduction

Variability in perception and action is a prominent topic in neuroscience (Ditzinger & Haken, 1989; Leopold et al., 2002; Lumer et al., 1998; Schmidt et al., 1979) and can be observed even when the external conditions, such as the sensory input or task goal, are consistent across experimental trials. It is known that trial-to-trial variability in motor function is inevitable even for well-trained experts, and similar arguments have been made regarding perception (Dhawale et al., 2017). Variability can be attributed to a variety of factors, of which the most fundamental is the variability of the nervous system’s response. Therefore, the effects of neuronal variability on behavioral variability have been widely discussed in the context of neuroscience.

Neuronal variability is evident at the neuronal level from the observation of single-cell spikes (Mainen & Sejnowski, 1995) to electroencephalograms (EEG) (Arazi, Gonen-Yaacovi, et al., 2017; Arieli et al., 1996; Garrett et al., 2011) and can have various causes (Dinstein et al., 2015; Faisal et al., 2008). For instance, random fluctuations in voltage- or ligand-gated ion channels can cause membrane potential changes, and the resulting electrical noise can affect neuronal responses even in the absence of synaptic inputs (Faisal et al., 2008; John et al., 2000). Such stochastic noise is one of the factors that cause variability in nervous system responses, not only at rest but also during stimulus-evoked activity. Furthermore, deterministic fluctuations due to the nonlinearity of the nervous system also cause variability in neural activity. From a nonlinear dynamical systems theory viewpoint, it is natural that the neural system behaves inconsistently due to its complex connections. Nonlinear dynamical systems often exhibit complex and sensitive dependencies on initial conditions, meaning that even small differences in the starting state of a system can lead to significant differences in its behavior over time. This property holds for both the entire network and the microscopic level of the brain; neurons are known to perform highly nonlinear operations, and even small biochemical and electrochemical fluctuations can significantly alter whole-cell responses (Faisal et al., 2008).

Nevertheless, despite these inherent inconsistencies in the neural system, our perceptual and motor experiences are relatively consistent. The same object is perceived as such across time and space, and the body does not move unexpectedly depending on the situation. This raises questions about how the brain, a system with both deterministic and stochastic fluctuations, achieves consistent experiences, and how variabilities affect neuronal information processing (Dinstein et al., 2015).

Recent related studies have suggested that it is important for a healthy brain to show high variability in the resting state (with no input) and high consistency in its response to stimuli (Arazi, Censor, et al., 2017; Garrett et al., 2011; Schurger et al., 2015). For instance, it has been reported that younger and healthier individuals exhibit a greater degree of variability in brain activity and that increased variability correlates with better performance in cognitive tasks (Garrett et al., 2011). Other studies have reported that neural variability quenching, i.e., large fluctuations in brain activity during the resting state and small trial-to-trial variability during specific tasks, are important for task performance (Arazi, Censor, et al., 2017; Arazi, Gonen-Yaacovi, et al., 2017; Daniel & Dinstein, 2021; He, 2013; Schurger et al., 2015). It has also been found that individuals with disabilities, such as those in a vegetative state, have a weaker degree of variability quenching than healthy individuals (Schurger et al., 2015). Additionally, several fMRI and EEG studies have reported a higher degree of neural variability in patients with psychiatric disorders (e.g., autism spectrum disorder and attention deficit hyperactivity disorder) and during certain perceptual (visual, auditory, or tactile) tasks compared with controls (Dinstein et al., 2012; Gonen-Yaacovi et al., 2016; Haigh et al., 2015; Hornickel & Kraus, 2013; Weinger et al., 2014).

In nonlinear dynamical systems, the property of identical responses to the same time-varying input regardless of differences in the initial state of the system is called “consistency” and is widely investigated in various nonlinear systems, such as laser systems (Uchida et al., 2004), the Lorenz model (Uchida et al., 2008), and the human brain (Kitajo et al., 2020). These considerable results and the perspective of nonlinear dynamical systems theory suggest that the brain benefits from a delicate balance between more complex dynamics in the absence of input and more consistent responses in the presence of input.

Neural consistency of the response to stimuli is not a common characteristic of the system but depends on the pattern of input received. In a pioneering study of trial-to-trial variability in neural activity to the same input, neurons in rat neocortical slices showed different consistencies depending on the temporal pattern of input at the level of single-cell responses (Mainen & Sejnowski, 1995). It has also been reported in macroscopic experiments that the brain shows selective, highly consistent responses to specific time-varying input patterns relative to other patterns (Andrillon et al., 2017; Kang et al., 2021; Luo et al., 2013). Additionally, it has been reported that the consistency of the nervous system is related to the consistency of perception and other abilities (Hornickel & Kraus, 2013; Luo et al., 2013; Schurger et al., 2015; Shishikura et al., 2023). For instance, Schurger et al. reported that subjects can perceive presented stimuli that are at a level of the sensory threshold when neural activity is consistent across and within trials (Schurger et al. 2015). In addition, a previous study of our group showed that spatio-temporal consistency of EEG responses to repeatedly presented television commercials accounted for population preferences associated with the engagement and memory of viewers (Hoshi et al., 2023). These studies suggest that the degree of neural response consistency depends on the characteristics of time-varying stimuli and affects cognitive and perceptual processes.

Here, we propose the term "selective consistency" to describe the phenomenon where neural activity exhibits high consistency and low variability in response to specific stimuli or time-varying patterns, even if those stimuli are not cognitively distinguishable. Neurophysiological studies have shown that highly variable neural activity augments the learning process and decreases to a well-learned pattern in the late learning phase (Dhawale et al., 2017). Additionally, it has been reported that the trial-to-trial consistency of neural activity improves rapidly for sensory stimuli that are repeatedly presented (Luo et al., 2013), regardless of the subject’s consciousness, understanding of the experimental purpose, or attention (Andrillon et al., 2015, 2017; Kang et al., 2021). Furthermore, a recent study found that the precision with which single neurons fire did not improve with learning, suggesting that changes in neural consistency induced by learning do not occur at the level of single neurons but rather at a network level (Costa et al., 2017). These results suggest that improved selective consistency is implicitly achieved at the network level; however, the exact mechanism of this change is not clear.

Thus, we hypothesized that selective consistency is acquired in a self-organizing manner by plasticity within the nervous system, resulting in NRD learning. Self-organization is a phenomenon in which a system creates a structure with some kind of order as a result of individual autonomous behaviors and their interactions, even if the system cannot look at its entirety or adjust itself to suit its purpose. Changes in brain network connections also represent typical self-organization, which is mainly carried out by plasticity-induced renewal of synaptic connection strength between cells (Oja, 1982). In the present study, we propose a mechanism by which changes in selective consistency are achieved in a self-organizing manner by the plasticity of neural networks.

## 2. Noise repetition-detection task

This study aims to replicate a human behavioral task known as the noise repetition-detection (NRD) task (Agus et al., 2010) using a neural network to elucidate the underlying mechanism of selective consistency. This task was chosen for three reasons. First, it uses specialized stimuli that are not encountered in everyday life, implying that the learning effect is confined to the experimental setting (Agus et al., 2010; Agus & Pressnitzer, 2021; Kang et al., 2017). Second, changes in selective consistency in this task have been observed not only in awake humans but also in anesthetized mice and sleeping humans, suggesting that the effect is independent of cognitive functions (Andrillon et al., 2015, 2017; Kang et al., 2021; Luo et al., 2013). Third, this learning effect is not modality-specific but widely observed in the nervous system (Kang et al., 2018).

### 2.1. Details of the NRD task

In the NRD task, listeners are presented with a white noise stimulus and asked to report whether the stimulus contains a repetition of noise tokens or not. In basic NRD tasks, subjects were presented with either a 1 s sample of white noise (noise condition, N) or two identical and seamlessly concatenated 0.5 s white noise tokens (repeated noise condition, RN). For both N and RN stimuli, white noise realizations of sequences differed from trial to trial. Sporadically, one particular exemplar for both N and RN sounds reoccurred, interspersed throughout each experimental block and referred to as the referenced repeated noise (RefRN) and referenced noise (RefN), respectively (Fig. 1). That is, for RefRN stimuli, not only was the same token of noise repeated (two times) within a trial, but also the same realization reoccurred over multiple trials. Listeners were asked to report whether the presented noise time series had a repetition of noise tokens (RN) or not (N) for each trial. Since the subjects were only required to describe the stimulus type (RN or N) and were not made aware of the presence of Ref stimuli in each block, the learning was implicit and unsupervised (Agus et al., 2010; Andrillon et al., 2015; Kang et al., 2017, 2018; Kumar et al., 2014).

**Fig. 1.**
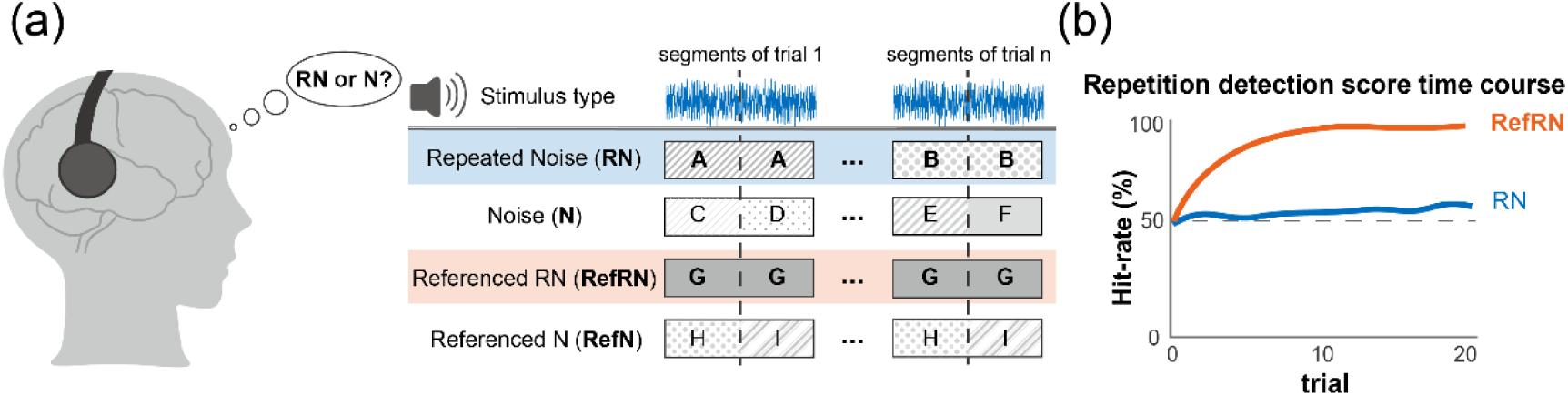
The noise repetition-detection task. (a) A conceptual diagram of the task and stimuli types. Participants listen to a 1 s white noise stimulus and answer whether the first and second 0.5 s segments are the same. There are four stimuli types, divided by the presence or absence of repetition within the stimulus (RN/N) and the presence or absence of repetition throughout the session (Referenced). As RN and RefRN have a repetitive structure, the correct answer is “Yes”, whereas for N and RefN, the correct answer is “No”. (b) The repetition detection score time course. The detection score for RefRN improves to a nearly perfect score immediately, whereas the score for RN remains around chance-level. As the participants do not know about the presence of the referenced stimuli, and the feedback for their answer is not given, this task can be regarded as implicit unsupervised perceptual learning.

Previous studies have shown that listeners can perceive time-series repetitions in white noise for stimuli that were previously encountered many times (typically five to ten) but cannot perceive repetitions of unfamiliar stimuli. The detection sensitivity and reaction time of listeners to RefRN gradually improved throughout a block, whereas RN performances remained at a chance level (Agus et al., 2010; Agus & Pressnitzer, 2013; Kumar et al., 2014; Luo et al., 2013). This gradual improvement in RefRN performance can be observed with a variety of sensory modalities and stimuli types, e.g., random visual and tactile pulse sequences (Kang et al., 2018).

### 2.2. Neural correlates of NRD task

Neurophysiological studies have suggested a relationship between NRD performance and the inter-trial consistency of neural activity. For instance, a study demonstrated that as learning proceeds, the N1-P2 complex auditory evoked potential of EEG in areas responsive to auditory stimuli becomes stronger and aligned (Andrillon et al., 2015). Event-related potentials are calculated by averaging hundreds of EEG waveforms for a specific event (in this case, white noise audio). Therefore, strong ERP components indicate that neural activity at that time is consistent within trials. Another study reported that the inter-trial phase coherence of the 3–8 Hz theta range of magnetoencephalogram after repeated exposure to stimuli was significantly higher than that for RN in the auditory cortex, even though they did not differ in early trials when learning was not sufficiently established (Luo et al., 2013). These changes in neural activities were also correlated with changes in behavioral performance (Andrillon et al., 2015). Furthermore, the enhancement of selective consistency can also be observed in neural activities in sleeping humans and anesthetized mice simply through stimulus presentation (Andrillon et al., 2017; Kang et al., 2021). Therefore, it was suggested that perceptual learning in the NRD task is associated with enhanced selective consistency, which occurs implicitly and distinctly from any cognitive function.

## 3. Methods

The model we used is based on the echo state network (ESN) (Jaeger, 2001) and has a plastic reservoir throughout unsupervised learning involving plasticity rules (Fig. 2). ESNs are artificial recurrent neural network models, which are characterized by feedback (“recurrent”) loops in their synaptic connection pathways. They can maintain ongoing activation even in the absence of input and thus exhibit spontaneous activity and dynamic memory. Biological neural networks are also typically recurrent networks. Therefore, ESNs have attracted attention as computational models of the information processing patterns of the brain (Enel et al., 2016; Jaeger, 2001; Lukoševičius & Jaeger, 2009). The classic ESN model has an input layer, a hidden recurrent layer, which is called a reservoir, and an output layer. The output of the reservoir is adjusted through the readout weights to match the target; however, there is no direct training performed over the reservoir, and it remains algorithms to work is that the underlying reservoir possesses the echo state property (ESP), which can be regarded as a particular shape of consistency (Boedecker et al., 2012; Gallicchio, 2018; Jaeger, 2001). When a drive subsystem and a response subsystem are represented by the states 𝒖 and 𝒙, respectively, the response subsystem given the time-series input from the drive subsystem converges to the state where 𝒙 is represented by 𝒙 = Φ(𝒖) and draws a repeatable trajectory for randomly connected (Jaeger, 2001). At this point, a crucial enabling precondition for ESN learning the same time-series input. Jaeger called such a trajectory a transient attractor and the property of converging to this trajectory an ESP (Jaeger, 2001). In simple terms, the ESP is a condition of asymptotic state convergence of the reservoir network under the influence of a driving input.

**Fig. 2.**
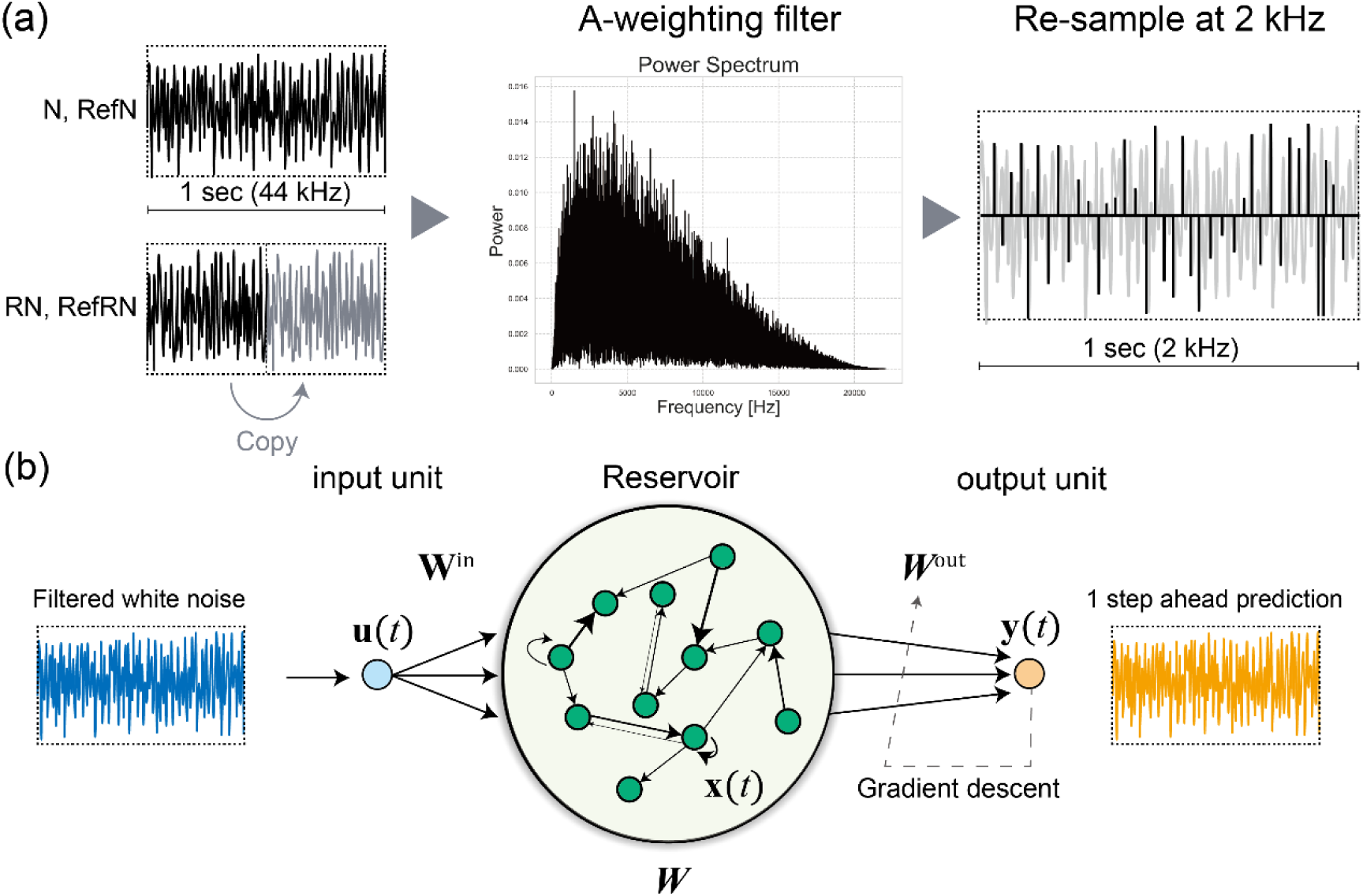
The model descriptions. (a) The process of input signal formation. There are four stimuli types: N and RefN stimuli consist of 1 s of white noise, whereas RN and RefRN stimuli consist of 0.5 s repetitions of white noise. The sampling frequency is 44 kHz. Subsequently, each time series is passed through an A-weighting filter, which reflects human auditory characteristics, peaking around 3,000 Hz, with high frequencies attenuated. After filter adaptation, each stimulus was resampled at 2,000 Hz to reduce computational costs. (b) An overview of the model. The resampled time series are presented to the neuron in the input layer as stimuli. 𝑾, the reservoir weights matrix is dynamic and maintained by Oja’s Hebbian plasticity rule. 𝑾^out^, the weights between the reservoir and neurons in the output layer are optimized using the gradient descent method. The model’s output target is one step ahead of prediction of the input time series.

In this study, we tested whether the ESN can acquire selective consistency (i.e., ESP depending on the input pattern) to a particular white noise realization via inducing plasticity into the reservoir. The ESP is connected to the algebraic properties of the reservoir weight matrix (Jaeger, 2001). Consequently, alterations to the reservoir weight matrix affect the reservoir’s consistency (ESP). To explore whether synaptic plasticity in the reservoir enables it to obtain selective consistency for an identical input, we incorporated Oja’s Hebbian rule into the reservoir (Oja, 1982). Note that changes to the reservoir weight matrix are implemented solely via input-dependent and unsupervised plasticity, rather than through processes such as prediction error minimization. This is a crucial point for modeling the acquisition of selective consistency in the brain. Because NRD learning can be observed even in the absence of task demands or awareness, the learning mechanism must be given without any supervised optimizations. The primary simulation concept is as follows: if ESNs with plasticity exhibit greater consistency to the Ref stimuli than to non-Ref stimuli after exposure to the stimulus set in the NRD task, then self-organizing changes in the network may be a mechanism to acquire selective consistency in the neural system.

### 3.1. Input signals

As our aim in this study was to simulate neural dynamics in the NRD task (Agus et al., 2010), the stimuli were generated in the same way (Fig. 2). There were four stimuli types: N, RN, RefRN, and RefN. The N and RefN stimuli consisted of 1 s white noise at a sample rate of 44 kHz. The RN and RefRN stimuli were repeated noise consisting of a repeated identical noise segment concatenated twice. N and RN stimuli were generated anew for each trial. RefRN and RefN were generated in the same way as RN and N; however, their realizations were the same for all trials within each condition (i.e., RefRN and RefN) throughout an experimental block. The auditory signals were generated as follows. First, each stimulus type was generated for 1 s (44,000 points). For RN and RefRN, the first and second segments (22,000 points) were set as the same time series. Second, all stimuli were filtered through an A-weighting filter, which is the most commonly used simple human auditory filter (Fletcher & Munson, 1933). Finally, sound signals were resampled to 2,000 Hz to reduce computational costs. For model training, five realizations for each stimulus type (20 stimuli in total) were presented as training data. The stimulus order was pseudorandom, except for the final presentation, which was set as one of N stimuli to avoid RefRN stimuli from being the last.

### 3.2. Neural network descriptions

The input signal, 𝑢(𝑘), is presented to the reservoir through the input weight matrix 𝑾^𝑖𝑛^ from the input layer neuron following:

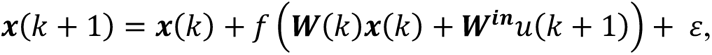

where 𝑥_𝑖_(𝑘), 𝑖 = 1, …, 𝑁 are the neural activations at time point 𝑘. ε replicates the internal gaussian noise and represents fluctuations of each neuron. 𝑓 is the activation function of the neurons and is defined as a hyperbolic tangent. 𝑾 ∈ ℝ^𝑁×𝑁^ is the synaptic weight matrix of the reservoir, which is fixed in the general ESN model. However, in this study, we arranged it to be dynamic and thus replaced it with 𝑾(𝑘). Several studies have demonstrated that the introduction of a variable called the "leaking rate" can enhance the computational performance of ESN (Jaeger, 2001). However, our research aims to examine changes in the system’s behavior induced by plasticity, rather than its computational performance; therefore, to simplify the problem, we did not introduce the leaking rate. The weight matrix, denoted by 𝑾^𝑖𝑛^ ∈ ℝ^𝑁×1^, connects an input neuron to the neurons in the reservoir and is generated as a set of uniformly distributed random numbers ranging from −1 to 1 and fixed throughout the simulation.

The number of reservoir neurons was set to 500, and their activation function was a hyperbolic tangent. The network was constructed as a sparse random network with a coupling density of d = 0.1. Non-zero elements of the weight matrix were defined as random numbers following a uniform distribution in the interval [−1,1]. To evaluate the impact of plasticity on reservoirs with varying degrees of consistency, we adjusted the spectral radius 𝜌(𝑾) of the reservoirs using the following modification:

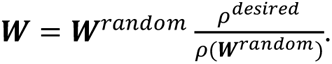

𝑾^𝑟𝑎𝑛𝑑𝑜𝑚^ is the weight matrix that was initiated randomly without regard for a spectral radius, and 𝜌^desired^ is the desired spectral radius. The spectral radius is an important hyperparameter that controls the connection strength in the reservoir. Specifically, it refers to the maximum absolute value of the eigenvalues of 𝑾. In general, if the activation function 𝑓 of the reservoir is tanh,

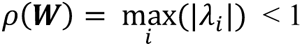

is a necessary condition for a reservoir to have ESP. Therefore, by adjusting the value of the spectral radius, we prepared various reservoirs near the edge of chaos (Legenstein & Maass, 2007) (the edge of having and lacking ESP).

To simulate the adaption to the input signals, Oja’s Hebbian rule was applied to the reservoir. This rule can be derived as a simple Hebbian plasticity rule that includes a forgetting factor to limit the explosion of the weight (Oja, 1982):

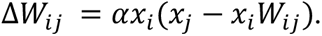

The synaptic learning rate parameter α was set to 10^−7^.

The output layer has a neuron that has connections to reservoir neurons with a readout weight matrix 𝑾^𝑜𝑢𝑡^ ∈ ℝ^1×𝑁^, and the output 𝑦 of this model is calculated as a linear sum of the reservoir neurons’ dynamics:

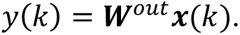

The computational task assigned to the model was to be one step ahead of predictions. Note that this study aimed to model how selective consistency is acquired implicitly. Neural selective consistency also occurs during sleep and in anesthetized mice, and thus we did not set the detection of noise repetition as the output task of the network in this simulation. The task assigned to the model was future predictions. This is based on the view of predictive coding, which assumes that the nervous system, especially the sensory system, works in a predictive manner (Denham & Winkler, 2020; Rao & Ballard, 1999). Therefore, we regarded the output signal as the neural dynamics in the network, which can be the basis of repetition detection, but not perception itself.

To maximize the performance for the time-series prediction, 𝑊^𝑜𝑢𝑡^ was optimized through the minibatch-based gradient descent (MBGD). The gradient descent method is an iterative first-order optimization algorithm used to find a local minimum/maximum of a given function. The error function that the gradient descent algorithm minimizes is written as follows:

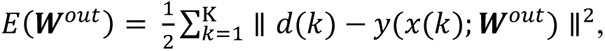

where 𝑑(𝑘) is the training data at time point 𝑘, and K is the total number of time points. The goal of the learning is to find the following:

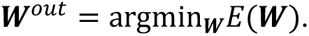

To solve the above problem, gradient ∇𝑬 is acquired. Then, the algorithm changes the variables 𝑾 for the negative direction of the gradient −∇𝑬 proportionally to the learning rate η:

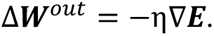

Hyperparameter η controls how rapidly the descent progresses, which is generally set at 0.01–0.00001. For our model, 20 statistically independent noisy inputs were given to reproduce the perceptual learning of humans, and we repeatedly applied the gradient descent method for each trial. This kind of pseudo-online gradient descent method is called “the minibatch gradient descent method” because the training data are divided into “minibatches”. The error of the minibatch number 𝑛, 𝐸_𝑛_, is calculated as follows:

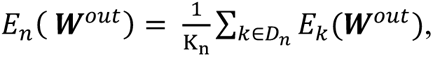

where 𝐸_𝑛_ reproduces the error of the minibatch number 𝑛, and K_n_ is the sample size of the minibatch data 𝐷_𝑛_. For normalization, the summation of error at each time point is divided by the sample size. The learning rate η was set to 0.01.

### 3.3. Simulation procedure

To investigate the effects of Hebbian plasticity, we trained multiple reservoirs with distinct spectral radii both with and without plasticity. The consistency of reservoir networks depends on their algebraic properties, particularly the spectral radius. A smaller spectral radius leads to consistent network behavior, whereas a larger one leads to chaos. We hypothesize that weak plasticity in the reservoir can fine-tune and make the reservoir have selective consistency to an identical input if the network is close to the edge of chaos. Therefore, we constructed reservoirs with spectral radii at values of 0.1–2.0 (see S1 Appendix for details). Subsequently, we exposed each reservoir to the stimuli of the NRD task and trained the reservoirs, both with and without plasticity, independently of each other. The training dataset and its order were set to be the same throughout the simulation.

### 3.4. Evaluation

We evaluated the changes in output time series, degree of selective consistency, and prediction error for each trained reservoir with 200 test runs. In the 200 test runs, N and RN stimuli used are different time series from those in the training runs, while RefN and RefRN used are the same time series as in the training runs. During the test runs, we halted the Hebbian rule and the gradient descent optimization of the output weights, and each trial was started with different initial values. This approach was chosen because in a nonlinear network, due to the network’s nonlinearity and ε—the internal Gaussian noise representing each neuron’s fluctuations—the network’s response exhibits different initial states and response trajectories in each trial. The reproducibility of these responses across trials can be the measure of consistency—the main topic of the present study.

#### 3.4.1. Consistency of outputs

Selective consistency was evaluated by Pearson correlations between 200 time points from the first and second halves of each output time series. The reason we don’t use the entire time series for both the first and second halves is that if the network exhibits ESP, it will eventually show a consistent response over time, while the duration of the transient period is different. The length of the window used for evaluation is based on the average duration of the transient period exhibited by the networks evaluated in this study—we set it at 200 points, which is one-fifth the length of the repetitive segment (see Figure 3).

**Fig. 3.**
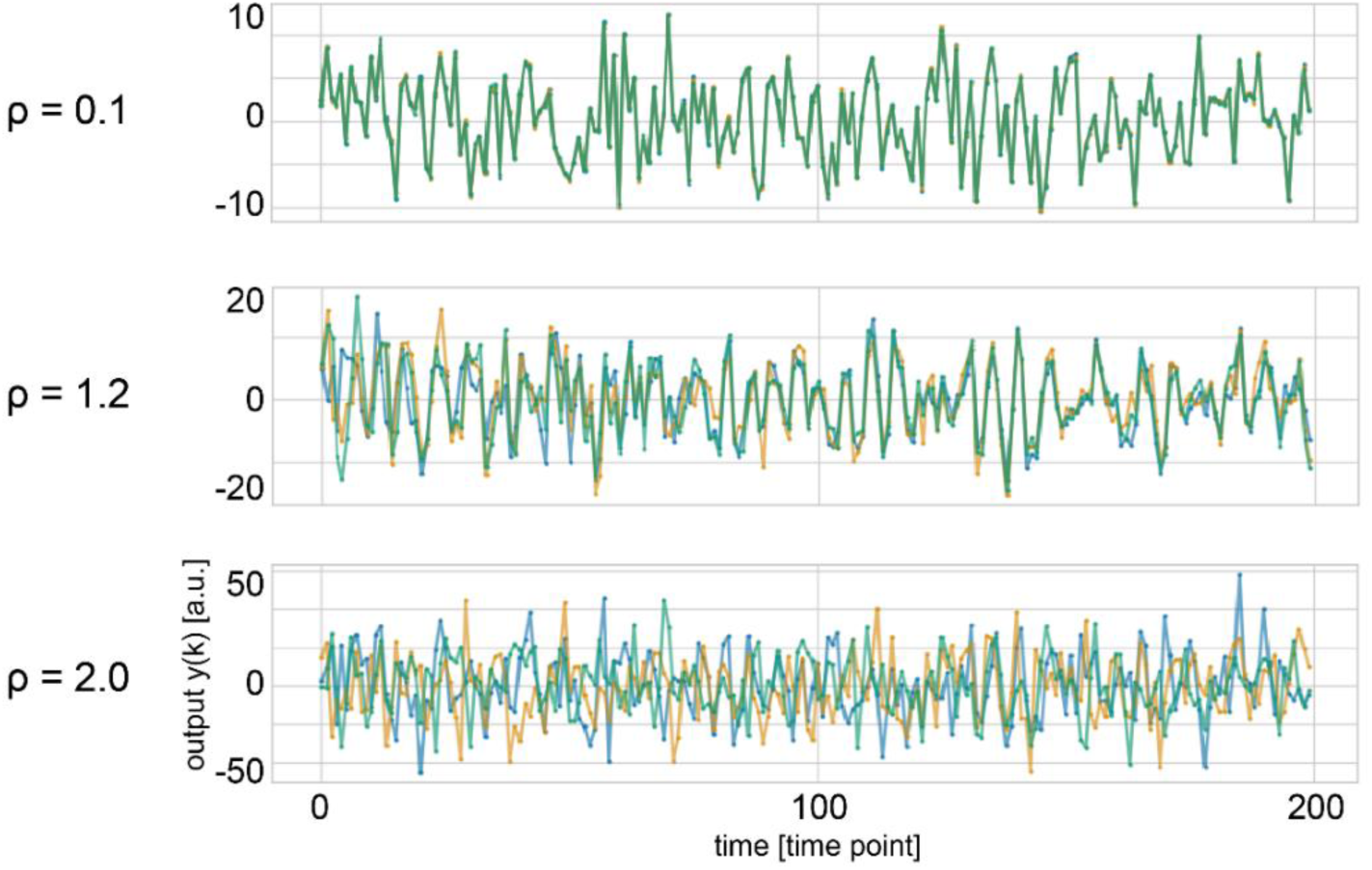
Representative examples of output time-series data. These are outputs for RefRN of Hebbian networks, although the similar tendency was observed for different network types and stimulus types. The results for three distinct spectral radii (ρ=0.1, 1.2, 2.0) are plotted separately. Each graph plots the overlaid output values on the vertical axis against the first 200 time points on the horizontal axis. The different colors correspond to different three output trials. Time is not plotted in its entirety, but rather, the first 200 points are magnified and plotted. When ρ is low, the lines converge following the transient period, indicating the identical response trajectory regardless of variations in the initial states. As ρ increases, the network no longer has ESP and behaves completely differently across distinct trials.

#### 3.4.2. Prediction error

we evaluated the accuracy of the time-series predictions learned in the ESN. Prediction accuracy of the time-series predictions were assessed using the root mean square error (RMSE) and normalized root mean square error (NRMSE) between the system output 𝑦(𝑘) and desired output 𝑑(k). The RMSE and NRMSE time-series with a spectral radius 𝜌, which is replicated as RMSE_𝜌_(𝑘) and NRMSE_𝜌_(𝑘), are defined as follows:

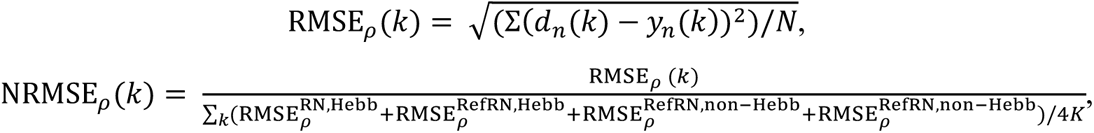

where 𝐾 represents the total number of time points, and 𝑛 stands for the number of trials. 𝑁 represents the total number of trials. RMSE^RN^ represents RMSE of RN of trial 𝑛 at one spectral radius. The definition of NRMSE is slightly different from the general normalization of RMSE, which is done by simply dividing RMSE by the mean of 𝑦(𝑘). This is because the output signal of the network tends to increase with larger spectral radii; thus, the normalization scale must be changed for each spectral radius so that this tendency does not affect the results. In addition, since the output time series is a white noise-like signal with zero mean, it was difficult to normalize by a simple mean, so we used RMSE for normalization. In this equation, normalization is performed by dividing by an averaged value of the RMSE of the four conditions: Hebbian vs. non-Hebbian and RefRN vs. RN stimuli.

#### 3.4.3. Statistical analysis

The statistical significance of selective consistency and the prediction error were tested with the nonparametric rank-order test based on the surrogate data technique (Lancaster et al., 2018). First, the difference between the evaluated selective consistency and prediction error of Hebbian and non-Hebbian paired distribution for each spectral radius and stimuli type were calculated, respectively. Next, we randomly shuffled these two pair-wise distributions and evaluated the difference between these two distributions to get a surrogate difference value. This was done 5000 times to obtain a surrogate difference distribution. The statistical significance level of the real data is determined by the percentile rank (p) of the difference obtained from the original data on this surrogate difference distribution. The results of statistical tests were adjusted for multiple comparisons using the Bonferroni method, considering the number of spectral radius levels and the combinations of stimulus types. The notation for percentile ranks (PR) used in this paper is as follows: (*; PR < 5%, p < 0.05, **; PR < 1%, p < 0.01, ***; PR < 0.5%, p < 0.005, ****; PR < 0.1%, p < 0.001).

## 4. Results

### 4.1. Output signals

We first investigated whether the output signal of the network exhibited the correct behavior as a predictive signal for a white noise time series. We plotted representative three randomly selected time series of the responses (Fig. 3). The behavior of the output signals was noise-like both for the Hebbian and non-Hebbian networks. Additionally, we found that even for the same network, the responses were different between distinct trials at the beginning of the output time series, which can be considered a transient period. This is attributed to the sensitivity of the network’s response to the initial state. This tendency became stronger as the spectral radius increased, and this is a typical characteristic of reservoir computing. The consistency of the system can be defined by the length of the transient period, and the initial state differences strongly influence the system outputs. If the system has strong consistency, its response will settle on the same trajectory after a short transient period. Therefore, we can say that both networks with and without plasticity show consistency in a range of relatively low spectral radii.

### 4.2. Selective consistency for RefRN

The inter-segment correlation analysis revealed that the plastic model obtained selective consistency for the RefRN stimulus, whereas the consistency for RN did not change. To evaluate the selective consistency for RefRN and RN, we compared the correlation between the first and second segments of output time series for RN and RefRN stimulus of both plastic and non-plastic models (Fig. 4). Significant differences in consistency were observed between the non-plastic and plastic models for RefRN, whereas no differences were observed for RN between both models.

**Fig. 4.**
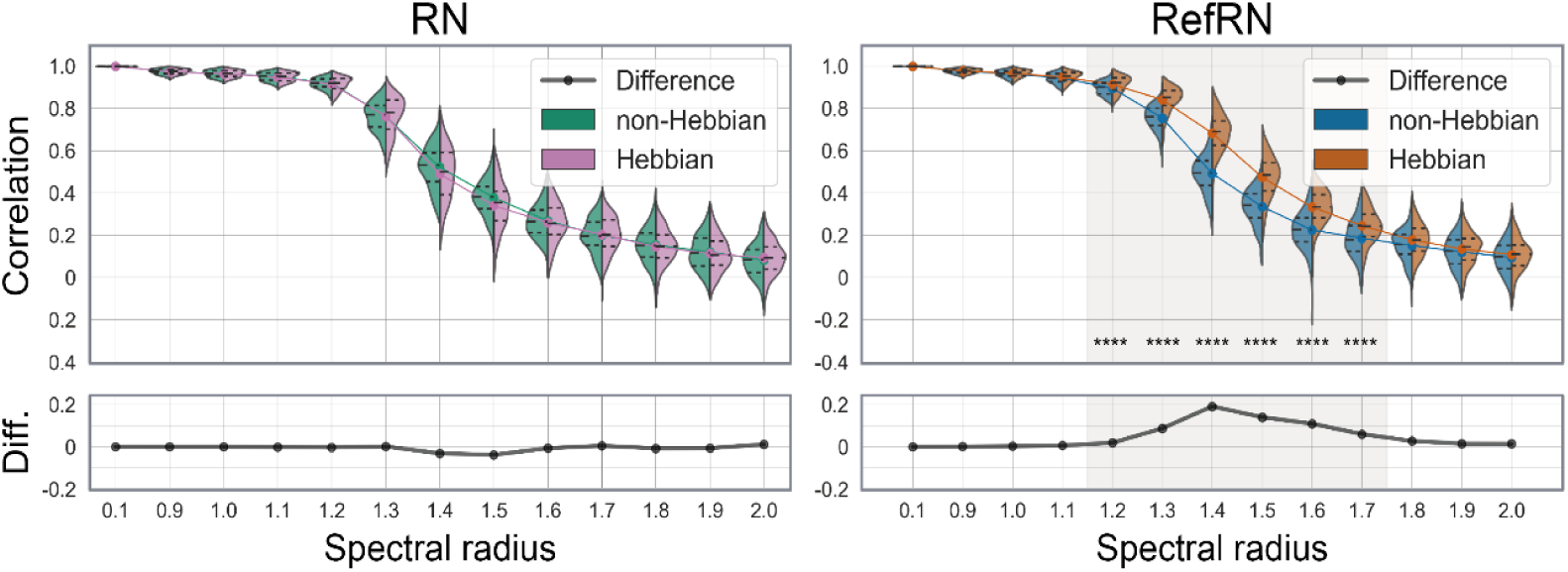
The evaluation of selective consistency. The consistency was evaluated by correlation between first and second segment time series for each test run with repeated noise (RN; left) and referenced repeated noise (RefRN; right). The violin plots show probability density distributions and interquartile ranges of Hebbian (right side; pink and orange) and non-Hebbian (left side; green and blue), respectively (****; PR < 0.01%, p < 0.001). The colored line plots connect the mean values for each condition. The black line in the bottom windows show the difference between Hebbian and non-Hebbian models. The horizontal axis represents the spectral radius of the evaluated networks.

Correlation decreased around where the spectral radius exceeded 1, regardless of the presence of plasticity. Generally, reservoir consistency is known to decrease as the spectral radius increases. However, the consistency of RefRN is enhanced by the presence of plasticity, especially in a regime between chaos and order. Conversely, the results of the RN showed no differences between plastic and non-plastic models. These indicate that the plasticity induced in the reservoir made the network obtain selective consistency for the repeatedly presented input signals. Furthermore, the prominence of selective consistency varied depending on the spectral radius. Selective consistency was not acquired in less complex or more complex reservoirs with a smaller or larger spectral radius than the critical area around 1.4 (Fig.5). The comparisons between the RefN and N conditions can be found in S2 Fig. Since the segments in these two conditions consist of different time series, the correlation coefficients were consistently near zero, regardless of the presence of plasticity or the spectral radius, showing no significant difference between the conditions.

**Fig. 5.**
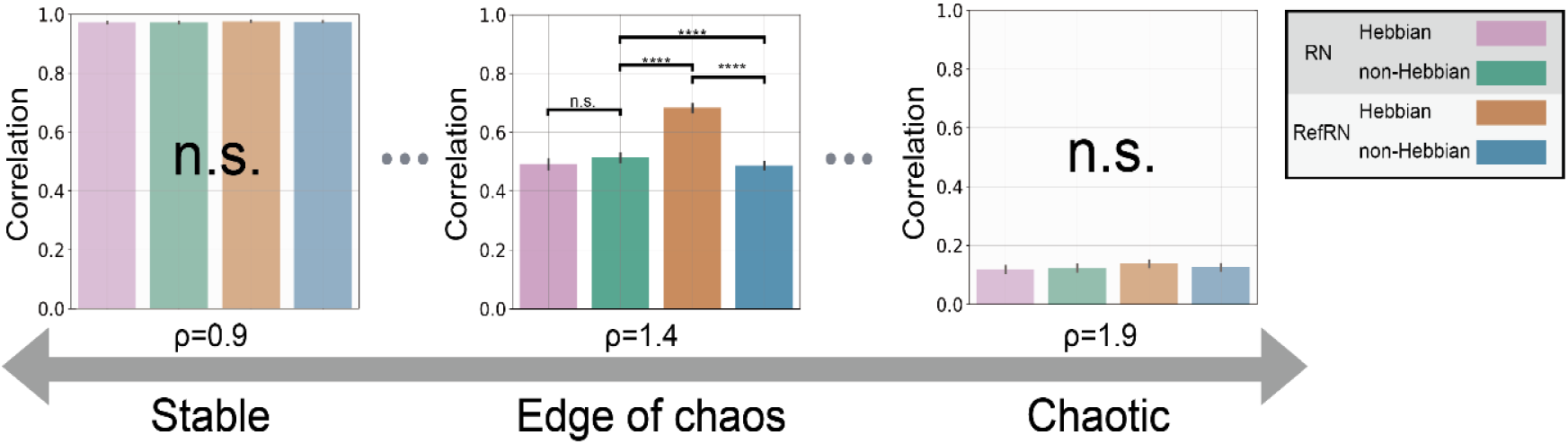
The edge of chaos and selective consistency. Each bin of the histograms shows averaged inter-segment correlation for four conditions (pink; RN of Hebbian network, green; RN of non-Hebbian network, orange; RefRN of Hebbian network, blue; RefRN of non-Hebbian network). The histograms represent results from networks with spectral radii of 0.9, 1.4, and 1.9, from left to right—which can be described as stable, edge of chaos, and chaotic, respectively (****; PR < 0.01%, p < 0.001). The error bars are representing 95 % confidence intervals. Notably, the edge of chaos region was chosen for its strong observation of selective consistency (see Fig.4). The statistical significance were tested with the nonparametric rank-order test based on the surrogate data model technique. It was found that in networks that are either stable or conversely chaotic, there was no difference between conditions, and differences were observed only in the edge of chaos region.

### 4.3. Selective consistency without optimization

As the consistency of the reservoir’s response depends on the spectral radius and input scale, the optimization of readout weights to obtain prediction accuracy may not play a crucial role in acquiring selective consistency. To test this, we also conducted the same analysis for the reservoir without optimizing readout weights 𝑊^𝑜𝑢𝑡^. Figure 6 shows the selective consistency without the optimization of readout weights. Similar to the results with the optimization process, significant differences in consistency were observed between the non-plastic and plastic models for RefRN, whereas no differences were observed for RN.

**Fig. 6.**
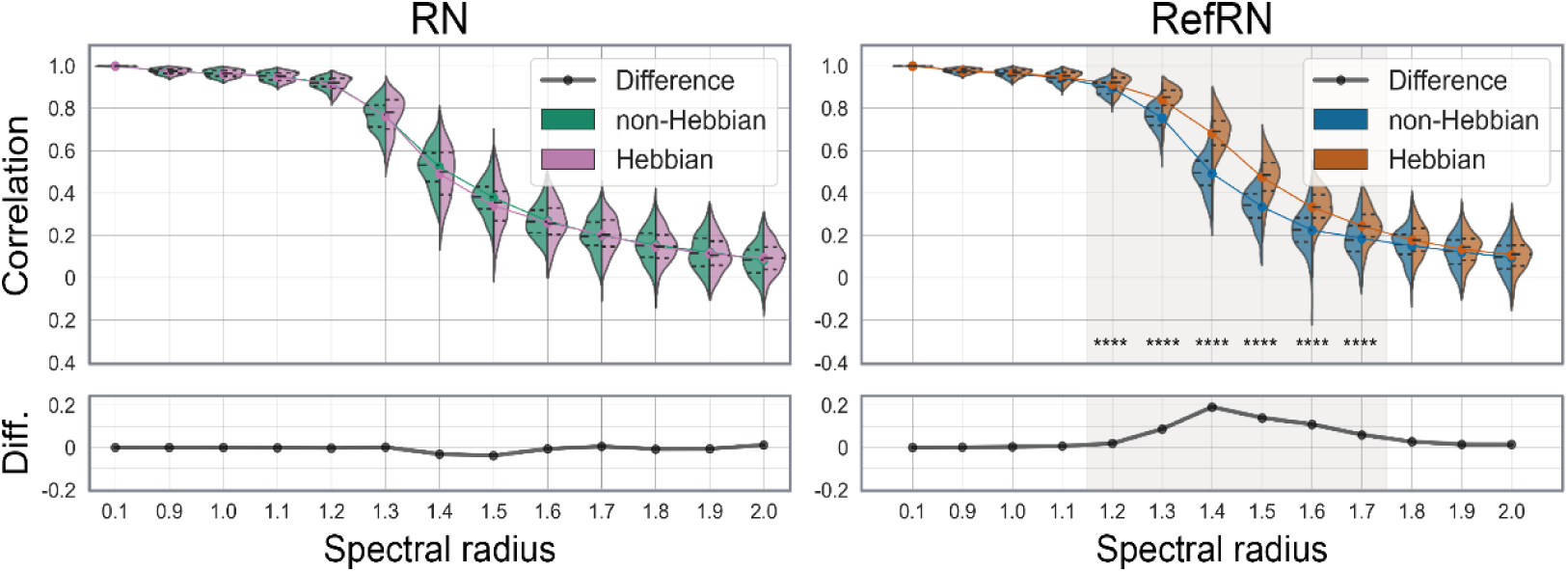
The evaluation of selective consistency without optimizing the readout weights. The figure styles are the same as Fig.4.The consistency was evaluated by the correlation between the first and second segment time series for each test run for repeated noise (RN; left) and referenced repeated noise (RefRN; right). The violin plots show probability density distributions and interquartile ranges of Hebbian (right side; pink and orange) and non-Hebbian (left side; green and blue) models, respectively (****; PR < 0.01%, p < 0.001). The colored line plots connect the mean values for each condition. The black lines in the bottom windows show the difference between Hebbian and non-Hebbian models. The horizontal axis represents the spectral radius of the evaluated networks.

### 4.4. Prediction error

The prediction error for each stimulus type was not affected by the presence or absence of plasticity spectral radius or stimulus type. As we verified above, the changes in the output layer’s connections due to optimization do not affect selective consistency. However, it is still unclear how the model’s temporal prediction accuracy changes with different spectral radii and the presence or absence of plasticity. Therefore, we finally examined the impact of changes in the accuracy of the time-series prediction. Figure 7 shows the prediction errors for RN and RefRN for varying spectral radii with and without plasticity. Although higher spectral radii resulted in higher prediction errors for both RN and RefRN, there were no differences in prediction error between RN and RefRN, regardless of the spectral radii or plasticity (Fig. 7a). This indicates that the prediction accuracy for repeatedly exposed stimuli (RefRN) is not selectively improved by introducing plasticity. Additionally, comparisons between NRMSE for each spectral radius did not show prominent selective minimization or maximization of prediction error for RefRN (Fig. 7b).

**Fig. 7.**
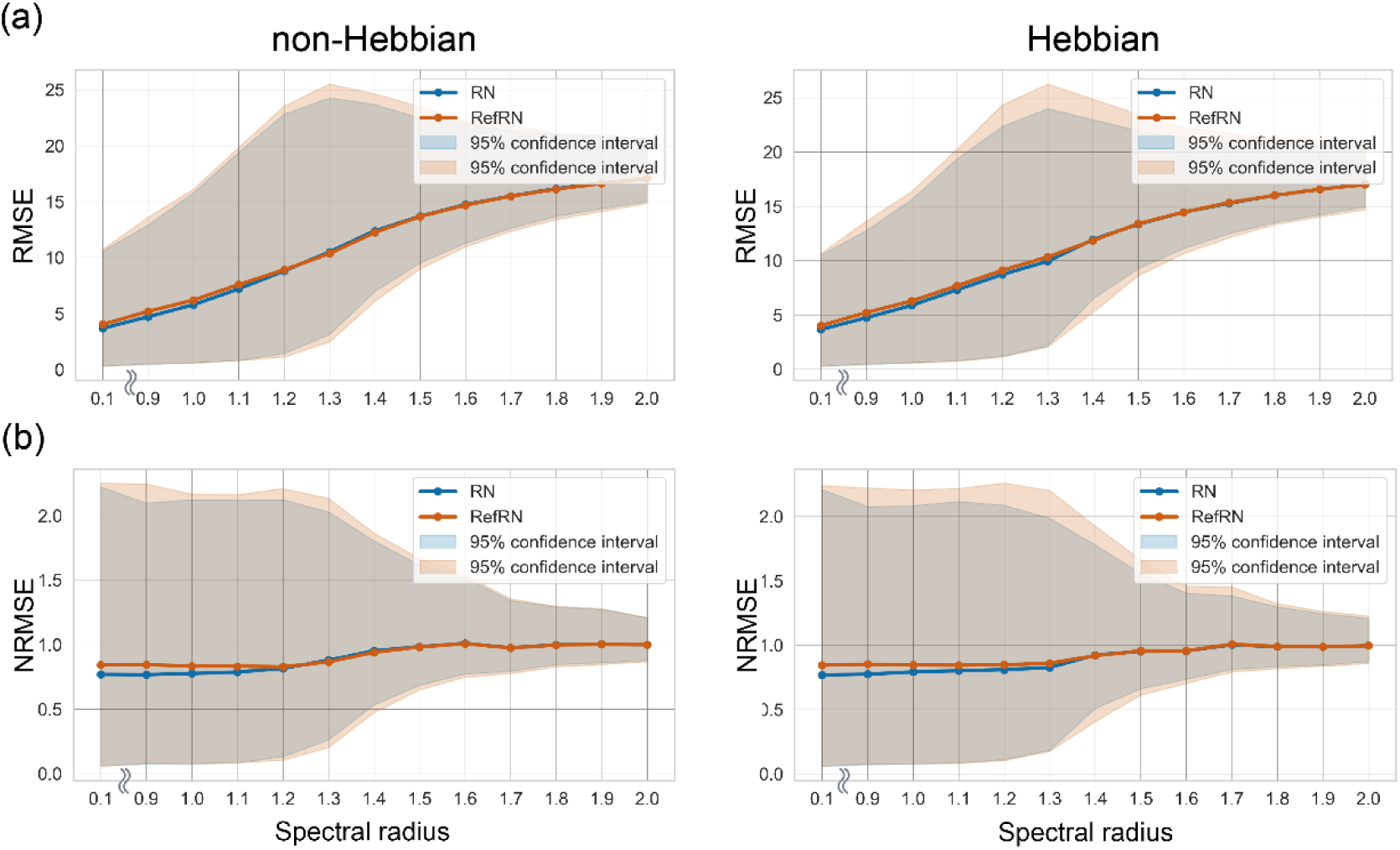
Prediction error with varying spectral radii. (a) Root mean squared error (RMSE) series and (b) normalized root mean square error (NRMSE) series. The plots represent the averaged RMSE or NRMSE between network prediction and the correct future time series. Non-plastic and plastic models are shown on the left and right, respectively. The results of RN and RefRN are represented by blue and orange lines, respectively. The 95% confidence intervals are depicted as filled areas.

## 5. Discussion

We have demonstrated that the network between order and disorder changes its composition in a self-organizing manner when a signal is given and that it acquires selective consistency for stimuli presented multiple times. In this study, we propose plasticity-driven self-organization of network connectivity as a mechanism by which neural networks enhance their response consistency to repeatedly exposed signals. A recent experimental study reported that the consistency of neural responses to the same input varies depending on the stimulus and is enhanced for repeated input stimuli in an implicit and unsupervised manner (Luo et al., 2013). We referred to this property as "selective consistency" and aimed to reveal its acquisition mechanism. It is not clear how neural networks achieve selective consistency; however, previous studies hypothesized that selective consistency could be acquired without any cognitive functions, such as attention and prediction (Andrillon et al., 2015), as it could be obtained in anesthetized mice (Kang et al., 2021) and sleeping human brains (Andrillon et al., 2017). Therefore, we validated whether the plastic network could obtain selective consistency using a reservoir-based plastic model. Several studies have previously attempted to introduce plasticity into reservoir computing (Morales et al., 2021; Schrauwen et al., 2008; Verleysen & Jochen, 2007; Yusoff et al., 2016). However, these studies were primarily based on a machine learning perspective, examining how the computational performance or memory capacity of the reservoir changes with the incorporation of plasticity— despite the crucial role of consistency in computation for recurrent networks. To the best of our knowledge, this study is the first attempt to investigate how the selective consistency exhibited by the reservoir changes due to self-organization.

Our simulation results reveal that the network could acquire selective consistency through plastic changes of recurrent networks at both inter-trial (S3 Fig) and inter-segment levels. Prediction errors increased as the spectral radius increased; however, this trend did not differ between stimulus conditions and did not depend on plasticity.

Furthermore, the changes in prediction error were linear within the range of observed spectral radii, whereas the change in selective consistency for RefRN was obtained only at the edge of the chaos region. This indicates that the acquisition of selective consistency occurred independently of the improvement in time-series prediction. One possible explanation for this contradiction is that the acquisition of selective consistency is achieved at the level of the reservoir neurons without the optimization of readout weights. In fact, the evaluation of selective consistency for a model without the optimization of readout weights yielded similar results to those of a model with optimization. This is in line with the hypothesis from previous experimental results that the acquisition of selective consistency occurs without the need for task presence or any top-down adjustments (Andrillon et al., 2015, 2017).

Although output optimization was not directly important for the acquisition of selective consistency, this does not mean that selective consistency is irrelevant to information processing, but rather that it plays an important role. As we argued before, the optimization process for the output weight does not play a crucial role in the acquisition of selective consistency. However, how the presence of selective consistency affects the optimization process warrants further discussion. In general, the computational performance of the reservoir-based model is known to depend on its consistency, and the network is required to have an ESP (Jaeger, 2001). This indicates that the improvement in the computational performance of the network (e.g., prediction accuracy) depends on the level of consistency. Therefore, in a situation in which the network has selective consistency for some input patterns, the optimization process is more affected by those reproducible patterns than by others with no reproducibility. This seems to be plausible for adapting to the environment and efficient information processing because the agent is not significantly affected by non-repeated noises that are likely to be ignorable. Thus, selective consistency may help the system reconcile the impact of statistically reproducible information with statistically non-reproducible information on model optimization based on its experience and consequently enable efficient information processing.

In addition, the prominence of selective consistency varied depending on the spectral radius and was strongest at the edge of chaos. The spectral radius adjusts the nonlinearity of the system, and in the case of an extremely small spectral radius, the system becomes linear and is not affected by the initial value. Therefore, a reservoir with a low spectral radius shows the same behavior for the same input regardless of the experiences. This is not in line with the behavioral experiment that shows consistency only for learned stimuli (Agus et al., 2010). Furthermore, considering biological facts related to the neural network structure (Allen et al., 2014; Friston, 1994) and numerous recent studies reporting the importance of variability in brain activity (Garrett et al., 2011), it becomes clear that overly linear responses are not suitable for neural system models. Conversely, if the spectral radius is extremely large, the system becomes chaotic and does not show consistency for learned and unlearned stimuli. Again, this is not suited for a neuronal activity model, which cannot adapt to the environment. From the results of the current study, it can be concluded that selective consistency through plasticity is not acquired when the spectral radius is too small or too large. This leads to the question as to why selective consistency is obtained only at a moderate spectral radius.

Generally, it is known that there is a critical point at which the behavior of the response significantly changes near a spectral radius of around 1.0 (Jaeger, 2001). Below this regime, the reservoir responds linearly, whereas values above result in chaotic behavior. Therefore, the computational performance of reservoir computing is maximized at a critical point, i.e., the edge of chaos. Recent fascinating research reported that some kind of recurrent neural network at the edge of chaos behaves chaotically in the absence of input signals but shows order in the presence of input signals and the computational performance of such networks is maximized(Haruna & Nakajima, 2019; Sussillo & Abbott, 2009). Our simulation also showed the best selective consistency for learned patterns near the edge of chaos. Therefore, our results may be interpreted as follows: the network that behaves chaotically in the absence of input can obtain ordered behavior selectively for a specific pattern. It is possible that consistency was obtained only for certain patterns due to the very weak level of plasticity introduced into the reservoir. This may have resulted in a very small change in the network, which, however, did not change the general consistency of the system.

In various contexts of different studies, it has been suggested that the brain exists at the boundary of order and disorder, the edge of chaos (Chua et al., 2012; Kumar et al., 2017). In addition to the brain, many other complex systems also exhibit a phenomenon called self-organized criticality, in which interactions between elements cause the entire system to transition to criticality (Dorogovtsev et al., 2008; Per et al., 1987). It has been suggested that the brain is a system that maintains a self-organized critical state, which it utilizes for efficient information processing (Plenz et al., 2021). In particular, the discussion of neural avalanches, which argues that the distribution of neural network activity follows a power law, serves as strong evidence that the nervous system exists in a self-organized criticality (Beggs & Plenz, 2003). In recent years, a model study using neural networks that reflect the physiological characteristics of the brain has reported conditions that realize both the edge of chaos and neuronal avalanches (Kuśmierz et al., 2020). As we discussed above, there has been active research on how the brain achieves a critical state and utilizes it for information processing. In our simulation results, self-organization was achieved through plasticity in the reservoirs, and it was confirmed that the learning effect was the strongest near the criticality. Furthermore, it is noteworthy that this area is not only realizing the strongest selectivity but also the only area that reproduces the experimental results of previous behavioral and neural activity measurements (Agus et al., 2010; Luo et al., 2013). Therefore, our results support the hypothesis that the brain utilizes maintaining a state close to criticality to adapt to the environment and to maintain a high information-processing ability.

Finally, the discussions we argued are not limited to a specific sensory modality but are a phenomenon that can occur in the nervous system in general. In this study, we developed simulations based on the human auditory behavioral task known as the NRD task. However, in practice, we simply gave white noise waveforms representing auditory stimuli to a network that mimics the nervous system and observed the changes that occurred due to the presence of internal synaptic plasticity. The same argument can be made for other sensory modalities as they are also input and processed as an electrical input waveform in the nervous system. Rapid perceptual learning similar to the NRD tasks has also been observed with visual and tactile stimuli (Gold et al., 2014; Kang et al., 2018). In addition, Kang and his colleagues have postulated that the performance in one modality can be predicted from the performance in other modalities (Kang et al., 2018). These considerations indicate that selective consistency itself and the ability or properties to acquire it are common and fundamental features of the entire brain.

It is important to consider the limitations of our research to properly contextualize our results. Despite its valuable findings, this study has certain limitations. First, our model does not fully elucidate the neural mechanisms of the NRD task. The aim of this study was to investigate the mechanism by which selective consistency is implicitly reinforced by experience, but not the mechanism by which the brain detects repetitions in auditory signals. Therefore, we did not make the model perform the NRD task explicitly. Researchers who are interested in the mechanism of repetition detection may need to investigate how the brain uses its selective consistency for selectively accurate information processing. There are several studies that are relevant to the mechanism of the NRD task (Andrillon et al., 2015; Masquelier, 2018). Again, although this study has utilized the NRD task, its aim was not to assess the task but to demonstrate that the consistency of network responses to the same input can be enhanced selectively, rapidly, and in a self-organizing manner.

Second, there are several models for reproducing plasticity in the nervous system other than the Hebbian rule, such as spike-timing dependent plasticity (STDP)(Gilson et al., 2011; Klampfl & Maass, 2013; Masquelier, 2018). Different results may be obtained by using another plasticity algorithm. However, a recent STDP-based learning model at the level of a single postsynaptic neuron showed that neurons progressively became selective to the repeating pattern unsupervised under certain STDP parameters, even if the pattern was embedded in strong background noise (Masquelier, 2018; Masquelier et al., 2008). Therefore, it can be said that the selective consistency presented in this study is not a narrowly model-dependent argument for plasticity.

Third, we confirmed that implicit enhancement of neural selective consistency can theoretically occur, as suggested by previous neural activity measurements (Luo et al., 2013). However, changes in consistency are possibly caused by some other mechanisms, and thus it cannot be claimed that all changes in consistency observed in the nervous system result from the manner we suggested. Finally, model simulations were used in this study, and thus it is still an open question whether such control of consistency exists in the actual nervous system. In a previous study that measured brain activity during the NRD task, the improved similarity of brain activity between trials was evaluated in terms of phase synchronization (Luo et al., 2013; Song & Luo, 2017). Our simulations suggest that not only inter-trial consistency but also inter-segment consistency for RefRN would selectively improve. As previous behavioral studies showed that the performance of repetition detection improved, we can assume that inter-segment selective consistency for RefRN also increases in the brain and is related to improved behavioral accuracy. To reveal the learning mechanism of the NRD task, it is necessary to evaluate whether biological neural activity also obtains selective consistency inter-segments. In this study, we regarded the constructed neural network as a particular form of a nonlinear dynamical system and investigated selective consistency. Given that the brain can be regarded as a similar nonlinear dynamical system, the analytical approach utilized in this study can be directly extended to the brain, enabling a direct comparison of our present investigation and empirical neural activity.

## 6. Conclusion

In the current study, we have demonstrated that a recurrent neural network with weak internal plasticity can acquire selective consistency, i.e., input-dependent consistency in its response. Despite many reports suggesting that the brain has overcome its inherent inconsistency to achieve selectively consistent activity and information processing, the neural basis of selective consistency remains unclear. Our computational simulations revealed that a reservoir-based plastic model could self-organize to acquire input-dependent consistency for frequently exposed stimuli. These findings indicate that a neural network near the edge of chaos can selectively optimize its response to frequently exposed inputs through fine self-organization due to weak plasticity. A learning model that accounts for dynamic changes in selective consistency may prove useful for understanding the brain’s efficient information processing mechanisms and for developing brain-like computers.

## Data availability

Data and code will be made available on request.

## Acknowledgments

This research was partially supported by JST Moonshot R&D Grant Number JPMJMS2292 and The Encouraging Grant for Graduate Students at NIPS

