## Supporting information for "Selective consistency of recurrent neural networks induced by plasticity as a mechanism of unsupervised perceptual learning"

| Parameter | Variable | Value |
|---|---|---|
| **Input dimension** | $N_u$ | 1 |
| **Reservoir dimension** | $N_x$ | $5 \times 10^2$ |
| **Reservoir connection density** | d | $10^{-1}$ |
| **Internal noise level in the reservoir** | ε | $10^{-1} \times N(0,1)$ |
| **Spectral radii** | $\rho$ | [0.1,0.9,1.0,1.1,1.2,1.3,1.4,1.5,1.6,1.7,1.8,1.9,2.0] |
| **Output dimension** | $N_y$ | 1 |
| **MBGD learning rate** | η | $10^{-1}$ |
| **Hebbian learning rate** | α | $10^{-7}$ |
| **Number of each stimulus type for a training dataset** | - | 5 |
| **Number of tests** | - | 200 |
| **Time points for transition** | - | 300 |

**S1 Appendix.** Experimental setting. For all simulations, we chose the following common parameters for the network constructions, MBGD, and Hebbian learning in this table.

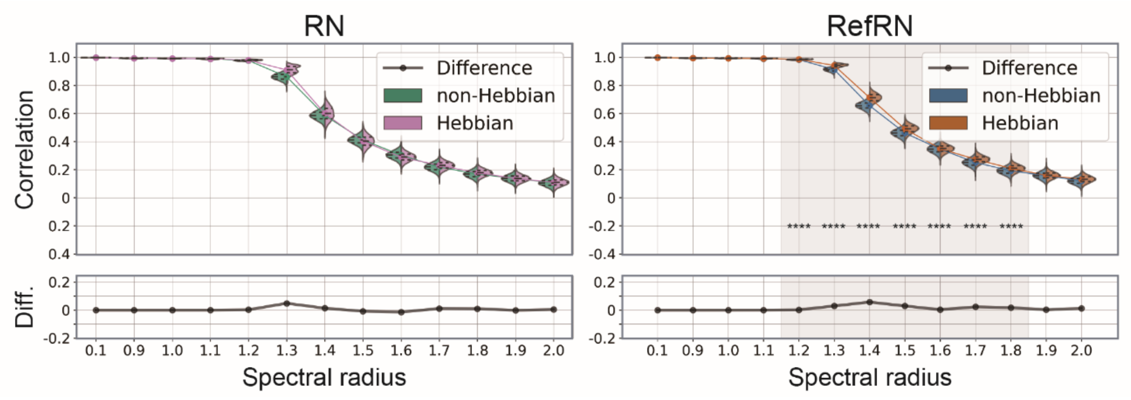


**S2 Fig**. The evaluation of selective consistency for RefN (right panel) and N (left panel) stimuli. The figure style is the same as Figure 4 and 6. The consistency was evaluated by the correlation between the first and second segment time series for each test run for repeated noise (RN; left) and referenced repeated noise (RefRN; right). The violin plots show probability density distributions and interquartile ranges of Hebbian (right side; pink and orange) and non-Hebbian (left side; green and blue) models, respectively (****; PR < 0.01%, p < 0.001). The colored line plots connect the mean values for each condition. The black lines in the bottom windows show the difference between Hebbian and non-Hebbian models. The horizontal axis represents the spectral radius of the evaluated networks.

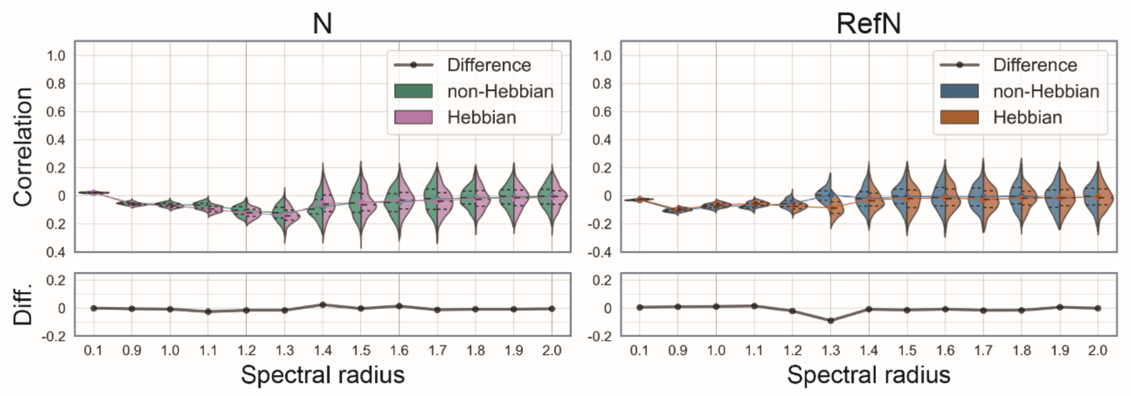


**S3 Fig**. The inter-trial level selective consistency for RefRN (right panel) and RN (left panel). The consistency was evaluated by the mean of the correlation between all time series.
The violin plots show probability density distributions and interquartile ranges of Hebbian (right side; pink and orange) and non-Hebbian (left side; green and blue) models, respectively (****; PR < 0.01%, p < 0.001). The colored line plots connect the mean values for each condition. The black lines in the bottom windows show the difference between Hebbian and non-Hebbian models. The horizontal axis represents the spectral radius of the evaluated networks.

Weaker but significant differences between conditions can be seen as same as the inter-segment level comparison.
